# A High Density Map for Navigating the Human Polycomb Complexome

**DOI:** 10.1101/059964

**Authors:** Simon Hauri, Federico Comoglio, Makiko Seimiya, Moritz Gerstung, Timo Glatter, Klaus Hansen, Ruedi Aebersold, Renato Paro, Matthias Gstaiger, Christian Beisel

**Affiliations:** Department of Biology, Institute of Molecular Systems Biology, ETH Zürich, Zürich, Switzerland; Competence Center Personalized Medicine UZH/ETH, Zürich, Switzerland; Department of Biosystems Science and Engineering, ETH Zürich, Basel, Switzerland; Biotech Research and Innovation Centre (BRIC) and Centre for Epigenetics, University of Copenhagen, Copenhagen, Denmark; Faculty of Science, University of Zürich, Zürich, Switzerland; Faculty of Sciences, University of Basel, Basel, Switzerland

## Abstract

Polycomb group (PcG) proteins are major determinants of gene silencing and epigenetic memory in higher eukaryotes. Here, we used a robust affinity purification mass spectrometry (AP-MS) approach to systematically map the human PcG protein interactome, uncovering an unprecedented breadth of PcG complexes. The obtained high density protein interaction data identified new modes of combinatorial PcG complex formation with proteins previously not associated with the PcG system, thus providing new insights into their molecular function and recruitment mechanisms to target genes. Importantly, we identified two human PR-DUB de-ubiquitination complexes, which comprise the O-linked N-acetylglucosamine transferase OGT1 and a number of transcription factors. By further mapping chromatin binding of PR-DUB components genome-wide, we conclude that the human PR-DUB and PRC1 complexes bind distinct sets of target genes and impact on different cellular processes in mammals.

## Introduction

Cell division requires faithful replication of the genome and restoration of specific chromatin states that form the basis of epigenetic memory^1^. Polycomb group (PcG) proteins – originally identified in *Drosophila melanogaster* as epigenetic regulators stably maintaining the repressed state of homeotic genes throughout development – are key players in this process. Numerous studies have now established a central role for PcG proteins in the dynamic control of hundreds of targets in metazoans, including genes affiliated to fundamental signaling pathways^2^. Hence, biological processes regulated by PcG proteins encompass cell differentiation, tissue regeneration and cancer cell growth^3–5^.

The PcG system is organized in multimeric repressive protein complexes containing distinct chromatin modifying activities, which impact on transcriptional regulation by modulating chromatin structures. In Drosophila, five distinct PcG complexes displaying different biochemical functions have been reported. The Polycomb Repressive Complex (PRC) 2 contains Enhancer of Zeste which trimethylates lysine 27 of histone H3 (H3K27me3)^6,7^ while the PRC1 subunit Polycomb provides binding specificity to H3K27me3 through its chromo-domain^8,9^. In addition, PRC1 also contains the dRing protein, which catalyzes the mono-ubiquitination of histone H2A on lysine 118 (H2AUb1), thereby blocking RNA polymerase II activity^10–12^. The Pho (Pleiohomeotic, Drosophila homolog of mammalian YY1) repressive complex PhoRC combines DNA-and histone tail binding specificities^13^, the PRC1-related dRing-associated factors complex dRAF contains the H3K36-specific histone demethylase dKDM2^14^ and the Polycomb repressive deubiquitinase (PR-DUB) targets H2AUb1^15^.

Although the core components of Drosophila PcG complexes seem rather fixed, we and others have shown that they can be co-purified with different sets of accessory proteins, thus increasing the diversity of the PcG system^13,16–18^. Epigenomic profiling revealed that distinct PcG complexes target largely overlapping gene sets in Drosophila and mechanistic details of PcG recruitment to target genes are beginning to emerge^15,19–22^.

In contrast, the mammalian PcG system is less well defined and appears to be significantly more complex. Each Drosophila PcG subunit has up to six human homologs, which combinatorially assemble in different complexes^23–26^. The six homologs of the Drosophila PRC1 core protein Psc, PCGF1-6, purify together with RING2, the homolog of dRing, in different complexes named PRC1.1-PRC1.6, and each of them associates with specific additional components^23–27^. These PRC1 complexes are further distinguished by the mutually exclusive presence of RYBP or a chromo-domain containing CBX protein. Five different CBX proteins displaying differential affinities for lysine-methylated histone H3 tails and RNA^28^ have been linked to PRC1. In contrast, the absence of a chromodomain within RYBP suggests that recruitment of CBX and RYBP containing PRC1 complexes might be mediated by H3K27 methylation or be independent of it, respectively. Indeed, recent work showed that the histone demethylase Kdm2b targets PRC1.1 via direct binding to unmethylated CpG islands^29–31^. Interestingly, incorporation of RING2 in optional PCGF complexes not only leads to differential recruitment to chromatin but also differentially regulates its enzymatic activity^23,29,31–34^.

Similarly to PRC1, the histone methyltransferase (HMT) activity of PRC2 is potentially modulated by accessory components such as the Polycomb-like homologs PHF1, PHF19 and MTF2^35–36^. Additional DNA binding interaction partners like JARD2 and AEBP2 might mediate recruitment of the complexes to chromatin^37–41^. However, whether the PRC2 core, consisting of EED, SUZ12 and EZH2, simultaneously interacts with all of these components or whether distinct complexes co-exist remains unknown. Moreover, mammalian PhoRC and PR-DUB have not been identified to date.

Understanding PcG-mediated epigenetic regulation in mammals requires a detailed understanding of the dynamic assembly of PcG complexes. A required step towards this goal is the exhaustive definition of the composition of individual PcG complexes including all accessory proteins, which likely convey distinct functional effects. Here we present the first systematic and comprehensive high-density map on the modular organization of the human PcG system using a sensitive double-affinity purification and mass-spectrometry (AP-MS) method^42–43^. The refined map of 1400 interactions and 490 proteins led to a considerable refinement of the human PRC1 and PRC2 network topology, including their relation with the heterochromatin silencing system and the identification of several novel interaction partners. Furthermore, we determined the composition of the human PR-DUB. We found that this highly diverse complex contains MBD proteins, FOXK transcription factors and OGT1, an O-linked N-acetylglucosamine (O-GlcNAC) transferase implicated in PcG silencing in Drosophila^44^. Finally, chromatin profiling of PR-DUB components and comparison with published chromatin maps of PcG proteins indicates that as opposed to Drosophila, PRC1 and PR-DUB regulate distinct sets of genes in human cells.

## Results and Discussion

### Systematic mapping of the human PcG interaction proteome

To investigate the human PcG protein interaction network, we applied a systematic proteomics approach, based on our previously reported AP-MS protocol in HEK293 cells^42^. The method employs Flp-In HEK293 stable cell lines expressing Strep-HA tag fusion proteins upon tetracycline induction (Fig. 1a). Initially, we selected 28 PcG proteins homologous to Drosophila core complex components and performed AP-MS experiments using these proteins as *primary baits* (Supplementary Fig. 1a). Then, based on the observed interaction data from this set, we chose 36 additional *secondary bait* proteins (Supplementary Fig. 1a, Supplementary Table 1). After double affinity purification, bait-associated proteins (preys) were identified by liquid chromatography tandem mass spectrometry (LC-MS/MS; Fig. 1a).

**Figure 1.**
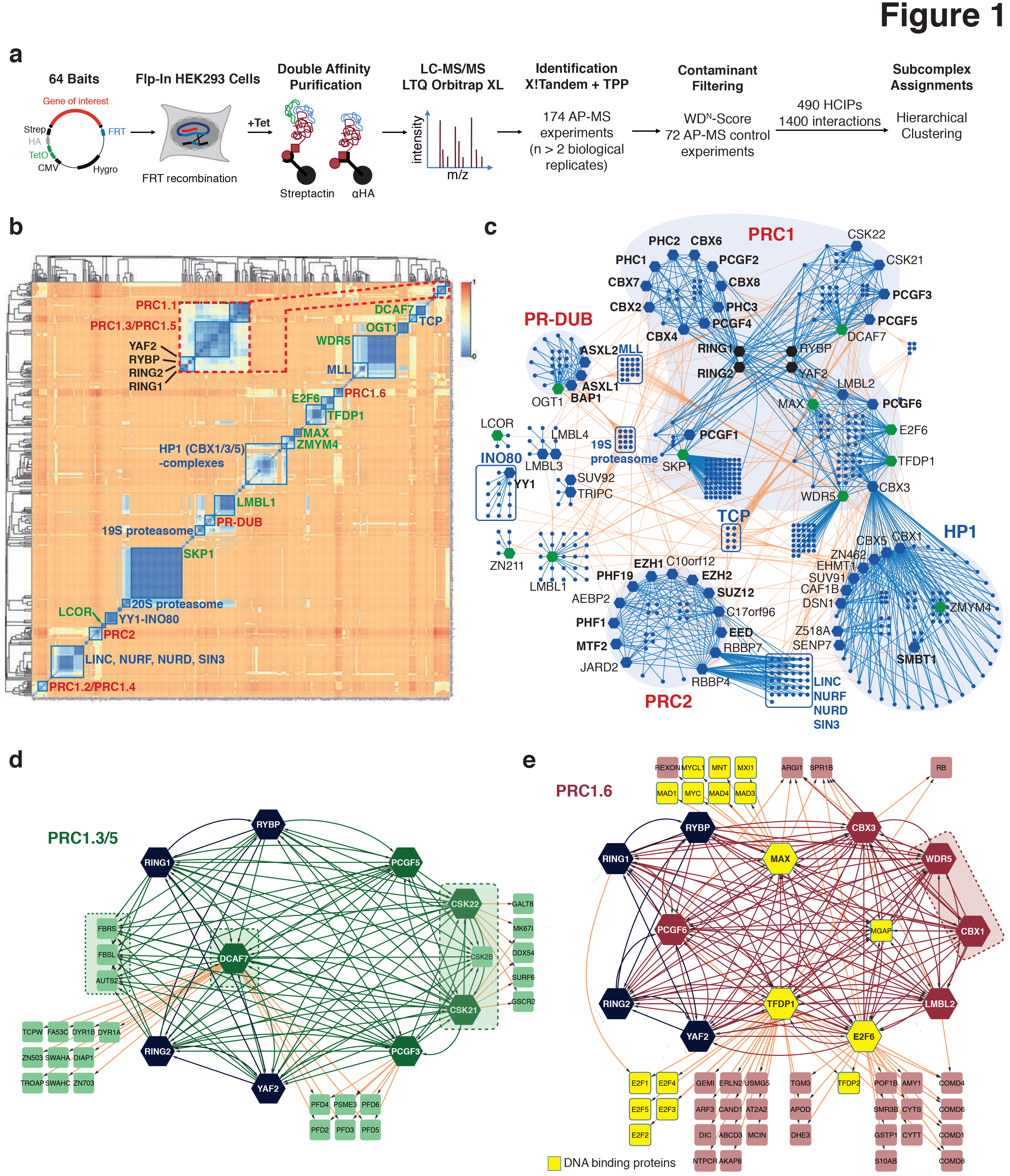
Systematic profiling of human Polycomb group (PcG) protein complexes. (a) Workflow for native protein complex purifications from Flp-In HEK293 T-REx cells. Open reading frames of 64 bait proteins were cloned into an expression vector containing a tetracycline inducible CMV promoter, Strep-HA fusion tag, and FRT sites. Proteins were affinity purified from whole cell extracts of isogenic cell lines, trypsinized and identified by tandem mass spectrometry on an LTQ Orbitrap XL. High confidence interaction proteins (HCIPs) were hierarchically clustered to infer protein complex compositions. (b) Hierarchical clustering of HCIPs. Clusters of PcG and non-PcG complexes are labeled in red and blue, respectively. The inset shows the location of PRC1.1, PRC1.3/PRC1.5 and the four core proteins RYBP/YAF2 and RING1/2. Clusters defined by single baits are indicated in green. Spearman’s rank correlation coefficient based dissimilarities are color coded as indicated (top right). (c) Protein-protein interaction network of clustered interaction data. Blue lines indicate interactions between proteins within the same cluster. Enlarged hexagon-shaped nodes correspond to the baits used in this study. (d-e) High-density interaction maps of PRC1.3/PRC1.5 (d) and PRC1.6 (e). New subunits are highlighted by dashed boxes. Hexagon shaped nodes represent baits; squares: identified HCIPs not used as baits in this study. Black nodes: common core subunits; yellow nodes: DNA binding proteins.

At least two biological replicates were measured for each bait protein, for a total of 174 AP-MS measurements. Proteins were identified using the X!Tandem search tool to match mass spectra to peptides, and the Trans-Proteomic Pipeline (TPP) to map peptides to proteins, at a false discovery rate (FDR) of less than 1%^45,46^. The resulting raw data set contained 930 proteins exhibiting 9856 candidate interactions.

To efficiently discriminate biologically relevant interaction partners from contaminant proteins, we devised a stringent filtering procedure based on both WDN-score^47^ and average enrichment over control purifications for each bait-prey pair. This filtering strategy retained 490 high confidence interacting proteins (HCIPs) encompassing 1400 (1193 unidirectional and 207 reciprocal) interactions. Our data set is characterized by an average of 21.9 HCIPs per bait protein, with 75% of interactions that have not yet been annotated in public databases (Supplementary Fig. 1b).

To evaluate the specificity and sensitivity of our AP-MS data, we considered the two bait proteins exhibiting the highest number of HCIPs, SKP1 (79 HCIPs) and WDR5 (73), and performed a cross-validation with literature-based reports. SKP1 serves as an adaptor for F-Box proteins and CUL1, and confers enzymatic specificity. Out of 79 HCIPs, our SKP1 purifications identified 42 F-Box proteins (Supplementary Fig. 1c). Furthermore, a previous AP-MS study investigating the interaction partners of WDR5^48^ identified a set of 21 proteins associating with this scaffold protein, which takes part in the assembly of several chromatin regulating complexes (reviewed in Migliori et al.^49^). Notably, while we were able to recall 76% of previously reported interaction partners, our experiments identified an additional set of 48 proteins (Supplementary Fig. 1d) co-purifying with WDR5 and encompassing MLL complexes, the NSL complex, the ADA2/GCN5/ADA3 transcription activator complex, mTORC2 components RICTOR and SIN1, and the Polycomb repressive complex PRC1.6 (Supplementary Fig. 1e).

### Hierarchical clustering assigns HCIPs to PcG complexes

To determine the topology of our protein interaction network, we performed hierarchical clustering of HCIPs using a rank-based correlation dissimilarity measure (see Supplementary Methods for details). Clustering revealed a modular organization built upon the three major PcG assemblies PRC1, PRC2 and PR-DUB, and HP1-associated complexes (Fig. 1b–c).

PRC1 represents the most elaborated and heterogeneous assembly, containing four groups of complexes defined by the six PCGF proteins: PRC1.1 (PCGF1), PRC1.2/PRC1.4 (PCGF2/4), PRC1.3/PRC1.5 (PCGF3/5) and PRC1.6 (PCGF6). Among these PRC1 assemblies, PRC1.6 further provides links to the heterochromatin control system via the HP1 chromobox proteins CBX1 and CBX3. Although analysis of the PRC1 topology has been recently reported in studies concentrating on specific subunits in various cellular systems^23,50–52^ our systematic high-density interaction data allowed us to further refine the composition of the PRC1 module. In the following discussion we focus on these novel findings regarding PRC1 organization, which is illustrated in Fig. 1b–e and Supplementary Fig. 3, and detailed in Supplementary Table 2.

All four PRC1 assemblies share a common core encompassing the E3 ubiquitin ligases RING1 and RING2, and – with exception of PRC1.2/PRC1.4 – RYBP and YAF2. Interestingly, PCGF2/4 also interact with RYBP and YAF2. As these proteins do not share any additional interaction partner besides RING1/2 (Supplementary Fig. 3b), RYBP/YAF-PCGF2/4-RING complexes might have limited functionality compared to other PRC1 complexes or correspond to transient products before specific canonical and non-canonical PRC1 holo complexes assemble. Furthermore, we did not detect any protein stably associating with all canonical PRC1 core members (RING1/2, PHC1-3, CBX2/4/6/7/8, PCGF2/4). However, we identified NUFP2 (Nuclear fragile X mental retardation interacting protein 2), a putative RNA binding protein exhibiting interactions with CBX2/6/7, PHC3 and PCGF4, as a new PRC1 interacting protein (Supplementary Fig. 3b).

The PRC2 complex is separated from both PRC1 and HP1 (Fig. 1c). The two characteristic histone binding proteins RBBP4 and RBBP7 not only belong to the PRC2 core along with SUZ12, EED and EZH1/2, but also partake in other protein complexes such as LINC, NURF, NURD and SIN3 (Supplementary Fig. 2a).

Finally, we identified the PcG complex PR-DUB defined by ASXL1/2 and BAP1 (Fig. 1c). Our clustering analysis also revealed complexes such as the TCP chaperonin and the proteasomal lid, that primarily consist of prey proteins (Fig. 1b). Of note, several proteins belonging to MLL complexes share interactions between PRC1.3/PRC1.5 (CSK21/22), PRC1.6 (WDR5) and PR-DUB (OGT1) (Fig. 1c). In contrast, interaction modules centered on LMBL1/3/4, SUV92 and TRIPC, LCOR, ZN211, and YY1 (the homolog of Drosophila Pho) are more disconnected and tend to be sparse (Fig. 1c, Fig. 2d and Supplementary Fig. 2b–Fig. c). Although YY1 interacts with all subunits of the INO80 chromatin remodeling complex, our AP-MS data does not unveil an equivalent of the Drosophila PhoRC complex (Supplementary Fig. 2b). However, except for PhoRC, we were able to reconstitute all mammalian equivalents of Drosophila PcG protein assemblies with unprecedented detail.

**Figure 2.**
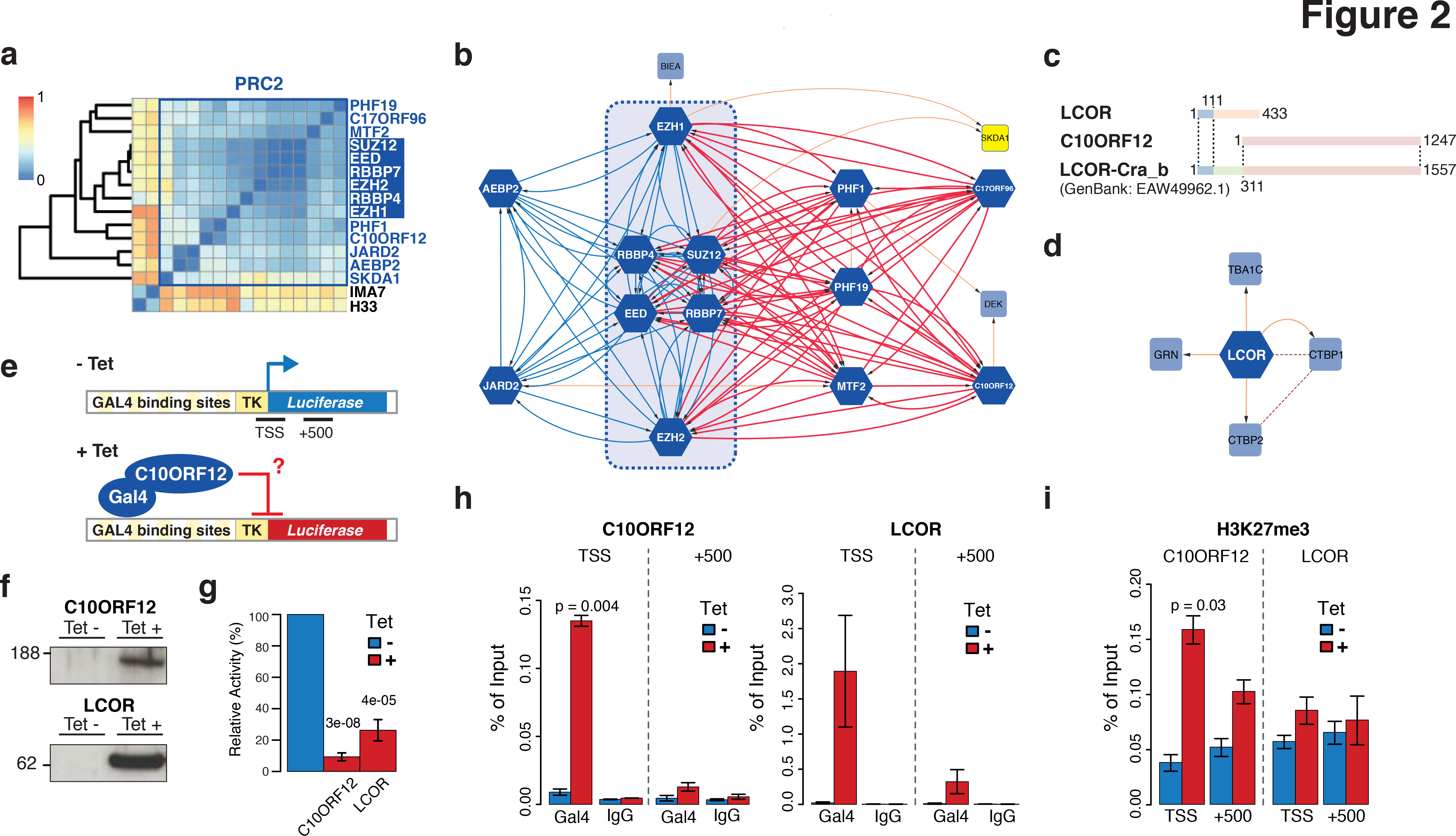
High-resolution interaction analysis unravels two structurally distinct classes of PRC2 complexes. (a) Excerpt of Figure 1b showing the PRC2 cluster. (b) Interaction map of PRC2 components. The PRC2 core is highlighted by a dashed box. Reciprocal interactions defining the two classes of PRC2 complexes are indicated in blue (PRC2.1) and red (PRC2.2) edges. Orange edges: non-reciprocal interactions. (c) Schematic representation of alternative protein isoforms of LCOR and C10ORF12. Numbers indicate amino acid positions. (d) LCOR interaction map. Orange edges, interactions defined in this study; dashed edges, published interactions. (e) Schematic representation of the employed luciferase reporter system. Amplicons (TSS, +500) used for ChIP-qPCR analysis are indicated. (f) Anti-Gal4 Western Blot showing the expression of Gal4-C10ORF12 and Gal4-LCOR upon tetracycline induction (g) Luciferase activity of tetracycline-induced Gal4-C10ORF12 and Gal4-LCOR expressing cells, normalized to uninduced cells. Values are mean?sd, p-values are from a two-sided t-test (n=7). (h) Anti-Gal4 ChIP-qPCR analysis showing localization of C10ORF12 and LCOR to the reporter TSS. (i) Anti-H3K27me3 ChIP-qPCR analysis at the reporter TSS upon C10ORF12 and LCOR expression. Values are mean±sd, p-values are from a two-sided t-test (n=3).

### WD40 domain proteins DCAF7 and WDR5 are central scaffolding proteins for PRC1.3/PRC1.5 and PRC1.6

The WD40 domain protein DCAF7 has been implicated in skin development and cell proliferation by interacting with DIAP1 and the dual-specificity tyrosine phosphorylation-regulated kinase DYR1A^53,54^. Intriguingly, DCAF7 co-purified with CBX4/6/8, RING1/2, RYBP/YAF2 and PCGF3/5/6, indicating that the protein is deeply embedded in the PRC1 module. As recent studies also reported interactions between DCAF7 and members of the canonical PRC1 complex, as well as PCGF3/5/6^26,27,55,56^, we performed DCAF7 purifications to test whether the protein is indeed a universal subunit of several different RING1/2-containing complexes.

Our DCAF7 AP-MS revealed reciprocal interactions with all bait proteins within a cluster centered on PCGF3 and 5 (Fig. 1d and Supplementary Fig. 3c), with no relation to the other PCGF complexes. Moreover, we identified DYR1A/B, DIAP1, the Zinc finger transcription factors (ZNFs) ZN503 and ZN703, and the ankyrin-repeat proteins SWAHA and SWAHC as an unrelated module interacting with DCAF7 (Fig. 1d). This result suggests that DCAF7 acts as a scaffold for several different protein complexes.

As for RING1/2, RYBP/YAF2 and PCGF3/5, DCAF7 interacts with the tetrameric casein kinase 2 (CSK2) and the three paralogs AUTS2, FBRS and FBSL. Therefore, to further refine the PRC1.3/PRC1.5 sub-network we performed AP-MS experiments using the catalytic casein kinase subunits CSK21 and CSK22. Our results confirmed the topology of the PCGF3/5-DCAF7 assemblies, and identify CSK2 and three uncharacterized proteins within the AUTS2 family as part of PRC1.3/PRC1.5 (Fig. 1d and Supplementary Fig. 3c).

The protein PCGF6 was initially purified together with the transcription factors E2F6, MAX, TFDP1, MGAP as well as RING1/2, YAF2, LMBL2, CBX3 and the HMTs EHMT1 and EHMT2, an assembly denoted as E2F6.com^57^. However, subsequent studies were unable to recover the entire (holo) E2F6.com^23,27,58,59^. Moreover, recent data suggest that PCGF6 and RING2 might interact with the WD40 domain protein WDR5^23^. We therefore decided to revisit the topology of the PCGF6-E2F6 network and to probe WDR5 connectivity by adding MAX, TFDP1, E2F6, LMBL2, CBX3, EHMT2 and WDR5 to our bait collection. Our AP-MS experiments unraveled a high-density network including reciprocal interactions between all but one (EHMT2) baits within this set (Fig. 1e and Supplementary Fig. 2c), thus demonstrating that the major PRC1.6 complex resembles E2F6.com. In addition, MGAP, MAX, TFDP1 and E2F6 purifications revealed a rich set of transcription factors that can heterodimerize with these proteins but that are not part of PRC1.6 as they did not connect to any other component thereof (Fig. 1e).

Recently, WDR5 was also reported to be part of the Non-Specific Lethal (NSL) complex and to form a trimeric complex with RBBP5 and ASH2L, which stimulates the H3K4-specific activity of the SET1 HMT family members SET1A, SET1B and MLL1-4^48,60–62^. Interestingly, while we recalled these interactions, we additionally detected reciprocal interactions of WDR5 with all PRC1.6 subunits, thus demonstrating that WDR5 is a universal component of activating and repressing chromatin modifying complexes.

Taken together, our results identify the WD40 domain proteins DCAF7 and WDR5 as subunits of PRC1.3/PRC1.5 and PRC1.6, respectively. Importantly, recent studies suggested that the diversity of PRC1 complexes might be specified by binding preferences of PCGF proteins, which are mediated by their RING finger-and WD40-associated Ubiquitin-Like (RAWUL) C-terminal domain^63,64^. For example, the PCGF1 and PCGF2/4 RAWUL domains selectively interact with BCOR/BCORL and PHC proteins, respectively^63^. Since no interaction partners of PCGF3/5 and PCGF6 RAWUL domains have been experimentally identified to date, and since WD40 domain-containing proteins often scaffold multisubunit complexes^49^, we propose that DCAF7 and WDR5 may serve as central scaffolding proteins for PRC1.3/PRC1.5 and PRC1.6.

### CBX1 partitions in several distinct heterochromatin complexes including PRC1.6

In contrast to previous studies, which reported CBX3 as the only heterochromatin protein within E2F6.com, we unexpectedly detected CBX1 in all our PRC1.6-related pull down experiments. To corroborate this finding, we performed AP-MS experiments with CBX1, using the constitutive heterochromatin protein CBX5 as control. Our results indicate that while CBX5 is disconnected from the PCGF6-E2F6 network, components therein interact with CBX1 (Fig. 1e and Supplementary Table 2). Furthermore, they validate interactions of EHMT2 with CBX1 and CBX3 and, to our surprise, separate EHMT2 and EHMT1 from PRC1.6, suggesting a separate complex containing CBX1/3, EHMT1/2, ZNF proteins, as well as the KRAB-ZNF interacting and co-repressor protein TIF1B (Supplementary Fig. 4a).

Since the PcG and heterochromatin silencing systems are functionally and molecularly related through PcG CBX2/4/6/7/8 and HP1 CBX1/3/5 proteins (reviewed in Beisel and Paro^65^), we further explored the CBX1/3/5 core of our network seeking for potential connections between these two systems. This survey led to a refined topology of CBX1/3/5-containing complexes and identified new interacting partners (Supplementary Fig. 4b–Fig. e). However, we did not detect additional connections to PcG proteins, suggesting limited direct cross-talk between protein components of the two silencing systems.

### The PRC2 core partitions into two different classes of complexes

While the functional core complex of PRC2 is composed of SUZ12, EED, RBBP4/7 and either EZH1 or EZH2, additional accessory proteins have been identified which may regulate the H3K27 HMT activity of the complex and its recruitment to chromatin^66–68^. However, how these proteins are organized within PRC2 or whether they assemble into independent PRC2 subcomplexes remains largely unresolved. To elucidate the topological organization of PRC2 complexes we performed AP-MS experiments using 14 reported PRC2-associated proteins (Supplementary Fig. 1a).

Hierarchical clustering analysis assigned all PRC2 baits to a single cluster exhibiting high intra-cluster correlations (Fig. 1b and Fig.2a) and forming a high-density interaction network (Fig. 2b). However, when reciprocal interactions were taken into account, our data revealed two fundamental alternative assemblies linked to the PRC2 core, the first defined by AEBP2 and JARD2 and the second by the mutually exclusive binding of one of the three Polycomb-like homologs (PCLs) PHF1, PHF19 and MTF2, respectively (Fig. 2b).

Taken together, our results identify two structurally distinct classes of PRC2 complexes. We therefore propose a novel nomenclature for PRC2, in which we refer to the two PRC2 wings as PRC2.1 (mutually exclusive interaction of PHF1, MTF2 or PHF19) and PRC2.2 (simultaneous interaction of AEBP2 and JARD2). AEBP2 and JARD2 can directly bind to DNA and have been implicated in the recruitment of PRC2 and modulation of its enzymatic activity^37–39,67^. Interestingly, depletion of JARD2 has only a mild effect on global H3K27 methylation levels, suggesting that PRC2.1 might be primarily responsible for maintaining H3K27me3 patterns genome-wide.

### C10ORF12 and C17ORF96 are mutually exclusive subunits of the Polycomb-like class of PRC2 complexes

Our purifications of the PRC2 core members and PCLs identified two largely uncharacterized proteins, C10ORF12/LCOR and C17ORF96, as PRC2 interactors (Fig. 2b). Both proteins have recently been shown to reciprocally interact and co-localize on chromatin with EZH2^66^, but their placement within the PRC2 topology and their functional role remained unknown.

Purifications of C17ORF96 confirmed all interactions with PCLs (Fig. 2b) and computational sequence analysis revealed that C17ORF96 is present in all vertebrate genomes. Interestingly, BLAST identified a single protein related to C17ORF96 in the human genome, the SKI/DAC domain containing protein 1 (SKDA1) (Supplementary Fig. 5a). SKDA1 belongs to the DACH family, which is defined by the presence of a SKI/SNO/DAC domain of about 100 amino acids, and is involved in various aspects of cell proliferation and differentiation^69,70^. However, C17ORF96 lacks the SKI/SNO/DAC domain and its homology to SKDA1 is restricted to the C-terminus (53% sequence identity within the last 60 amino acids) (Supplementary Fig. 5a–Fig. b), suggesting that this region encodes an hitherto uncharacterized protein domain. Interestingly, SKDA1 also interacts with EZH1 and SUZ12 (Fig. 2b), suggesting that this putative C-terminal domain mediates the interaction of C17ORF96 and SKDA1 with the PRC2 core.

Initial analysis of C10ORF12, the second uncharacterized protein highly connected to the PRC2 core, identified peptides that ambiguously mapped to two distinct UniProt proteins, LCOR and C10ORF12 (Supplementary Fig. 5c–e). These two proteins are encoded by the same genomic locus. Indeed, in contrast to the UniProt database, Genebank contains the LCOR-Cra_b (ligand-dependent co-repressor, isoform CRA_b, EAW49962.1) entry, where the N-terminal 111 amino acids of LCOR are fused to C10ORF12 and the two regions are separated by a 200 amino acid spacer (Fig. 2c). LCOR is a ligand-dependent co-repressor interacting via its N-terminal domain with nuclear hormone receptors in a complex including CTBP and a number of histone deacetylases^71,72^. While our AP-MS analysis yielded peptides of the LCOR N-terminus, C10ORF12 and the LCOR-CRA_b specific spacer (Supplementary Fig. 5c), peptides of the LCOR C-terminus were missing (Supplementary Fig. 5d), indicating that PRC2 interacts with LCOR-CRA_b and potentially with the shorter isoform C10ORF12. To test this possibility, we performed additional AP-MS experiments using LCOR, C10ORF12 and LCOR-CRA_b as baits. LCOR purified with its known interaction partners CTBP1 and CTBP2, while PRC2 components were absent in LCOR purifications (Fig. 2d and Supplementary Fig. 5e). In contrast, both LCOR-CRA_b and C10ORF12 reciprocally interact with all subunits of the PCL wing of PRC2 (Fig. 2b, Supplementary Fig. 5d–e).

To investigate the functional relevance of this finding, we employed a heterologous reporter system based on a stably integrated, constitutively active luciferase reporter gene responsive to upstream, promoter-proximal GAL4 DNA binding sites (Fig. 2e)^73^. We engineered cell lines containing tetracycline inducible GAL4-LCOR and GAL4-C10ORF12 expression constructs, respectively. Upon induction, both proteins accumulated in the nucleus and were recruited to the GAL4 motifs, resulting in strong repression of luciferase activity (Fig. 2f–h and Supplementary Fig. 5f). To assess whether the repressive activity of C10ORF12 is mediated by recruitment of PRC2 to the target promoter, we performed chromatin immunoprecipitation (ChIP) with an H3K27me3-specific antibody and analyzed the enrichment of luciferase promoter fragments via quantitative PCR. Upon tetracycline induction, we found that the transcription start site (TSS) of the luciferase gene was significantly trimethylated at H3K27 in the GAL4-C10ORF12 expressing cell line (Fig. 2i). In contrast, despite GAL4-LCOR was expressed at higher levels than GAL4-C10ORF12 (Fig. 2f) and exhibited a 10-20 fold increase in its binding to the reporter (Fig. 2h), no significant H3K27me3 enrichment was observed upon expression of this protein.

PCL proteins target PRC2 and positively regulate its enzymatic activity via their ability to bind methylated H3K36^36,74,75^. However, further experimental investigation will be required to elucidate the exact mechanism by which C17ORF96 and LCOR-CRA_b/C10ORF12 influence PRC2.1. An interesting possibility is that LCOR-CRA_b recruits PRC2.1 to nuclear hormone receptor binding sites upon ligand binding. This interaction, restricted to C10ORF12, leaves the N-terminus of LCOR free for ligand responsive interaction with nuclear hormone receptors.

### ASXL1 and ASXL2 define optional PR-DUB complexes containing OGT1 and FOXK transcription factors

The Drosophila PcG complex PR-DUB was identified as a heterodimer consisting of the deubiquitinase Calypso and the Asx protein^15^. However, the composition of its human counterpart remains elusive. Thus, we set out to systematically characterize this complex by performing purifications of BAP1, ASXL1 and ASXL2, the human homologs of the Drosophila PR-DUB components. Our AP-MS analysis revealed that BAP1 reciprocally interacts with both ASXL1 and ASXL2 (Fig. 3b). Interestingly, the two ASXL proteins do not interact with each other (Fig. 3b), suggesting the existence of two mutually exclusive PR-DUB complexes, which we called PR-DUB.1 and PR-DUB.2 depending on the ASXL partner of BAP1 being ASXL1 and ASXL2, respectively.

**Figure 3.**
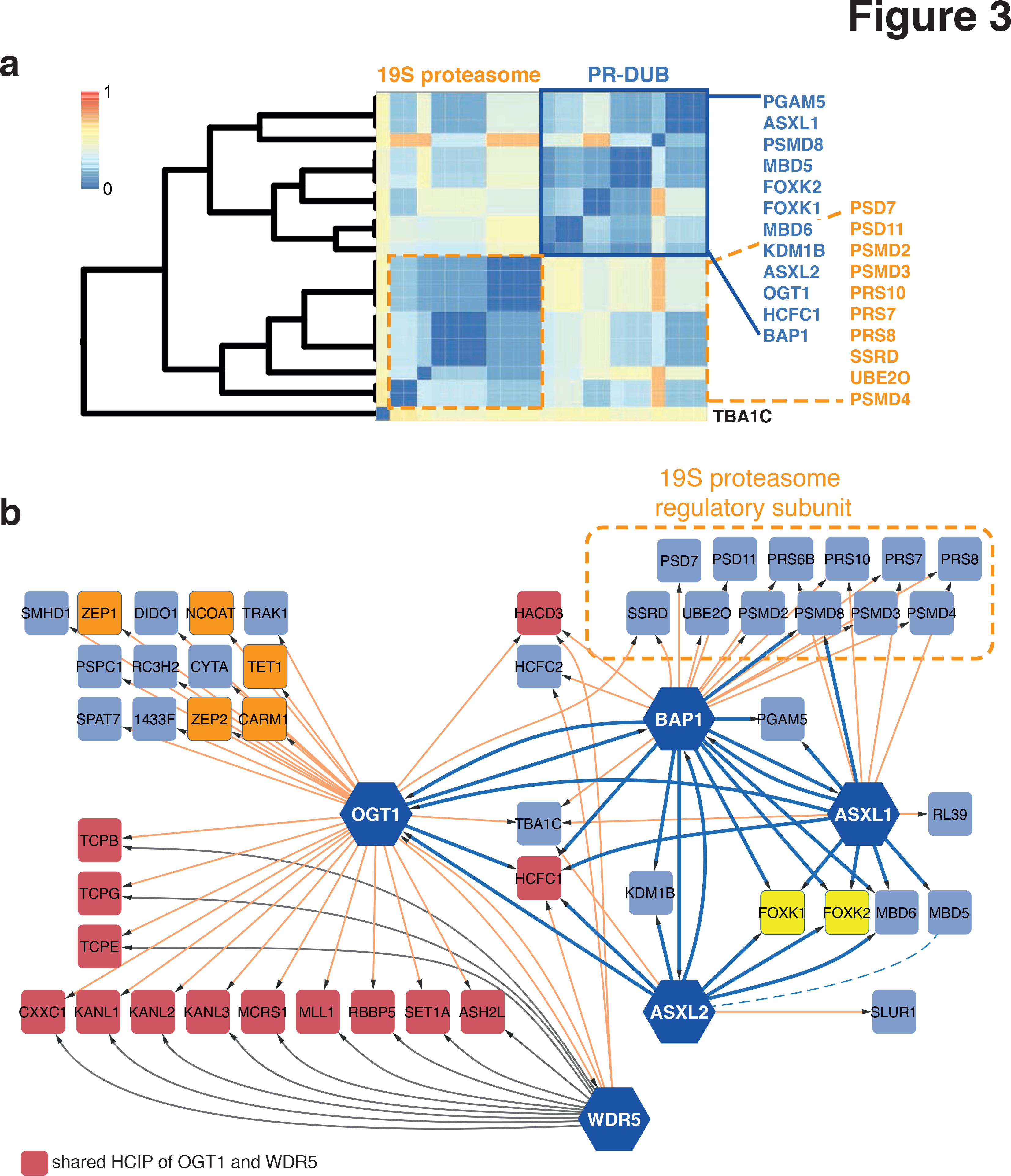
Human PR-DUB complexes contain OGT1 and FOXK transcription factors. (a) Excerpt of Figure 1b showing the PR-DUB and 19S proteasome clusters. (b) Topology of PR-DUB complexes. Interactions of bait proteins with proteins localized in PR-DUB cluster are indicated in blue. WDR5 shares many interacting proteins with OGT1 (indicated in red), which are predominantly MLL/SET complex associated proteins, and does not interact with BAP1, ASXL1 and 2. Hexagons: bait proteins; squares, identified HCIPs not used as baits in this study. Yellow: FOXK1 and 2.Orange nodes: OGT1 interactors. Dashed line: ASXL2-MBD5 interaction, which was detected but did not pass our stringent filtering criteria.

Both PR-DUB core components share a similar set of accessory proteins encompassing the transcription factors FOXK1 and FOXK2, the chromatin associated proteins MBD5 and MBD6, the transcriptional co-regulator HCFC1 and most notably OGT1 (Fig. 3b). A recent attempt to identify BAP1 interaction partners led to the identification of Asxl1, Asxl2, Ogt, Foxk1, Kdm1b and Hcf1 in mouse spleen tissue^76^. Our data provide support to these results and indicate a general, cell type independent assembly of mammalian PR-DUB complexes. Furthermore, our data clearly implicate OGT1 as member of mammalian PR-DUB complexes, an interaction which was not identified in the Drosophila PR-DUB complex purification^15^ although the Drosophila homolog Ogt was previously annotated as bona fide PcG protein^44^.

OGT1 is the only O-linked N-acetylglucosamine (O-GlcNAc) transferase in mammals. The enzyme catalyzes the addition of a single GlcNAc molecule to serine and threonine of many target proteins^77^. OGT1 enzymatic activity is required for mouse development and is essential for embryonic stem cell (ESC) viability^78^. In addition, the protein was found to interact with BAP1 and to localize to chromatin via its interaction with the 5-methylcytosine oxidase TET1^76, 78^. To further refine the connectivity of OGT1 within the PR-DUB network, we performed AP-MS experiments using OGT1 as bait.

This analysis validated the interaction between BAP1 and OGT1 and the interactions of OGT1 with TET1 and NCOAT (Fig. 3b), the O-GlcNAcase counteracting OGT1 activity^78, 79^. Moreover, our data identified a second set of OGT1-containing complexes involved in transcriptional regulation that did not co-purify with PR-DUB core subunits (Fig. 3b). These include the ZNFs ZEP1 and ZEP2, and the arginine-specific HMT CARM1. Furthermore, we identified OGT1 as subunit of WDR5 containing complexes. Indeed, OGT1 exhibits interactions with the NSL complex and with the SET1 HMT family activating complex WDR5/RBBP5/ASH2L, which is likely to mediate the interaction of OGT1 with MLL1 and SET1A (Fig. 3b). Although no interaction of OGT1 with FOXK1/2 and MBD5/6 was detected, these proteins co-cluster with PR-DUB core components and OGT1 is highly connected to the PR-DUB core (Fig. 3a–b).

These results suggest that OGT1/HCFC1 and FOXK/MBD proteins may form optional PR-DUB.1/PR-DUB.2 complexes. Conversely, OGT1 interactions with FOXK and MBD proteins could be transient and hence difficult to pinpoint by OGT1 affinity purification.

### Genomics profiling of the FOXK1-containing PR-DUB.1

A functional interaction of OGT1 with FOXK transcription factors within the same PR-DUB complex would require their colocalization at genomic target sites. To test this hypothesis, we examined the genome-wide distribution of O-GlcNAc, a proxy for catalytically active OGT1, ASXL1 and FOXK1 by performing ChIP-seq in HEK293 cells (Supplementary Fig. 6a).

By pairwise analysis of overlapping peak regions we found 41% and 55% of FOXK1 peaks co-localizing with O-GlcNAc and ASXL1, respectively, while 69% of O-GlcNAc peaks were co-occupied by ASXL1 (Fig. 4a). In total, we identified 2703 genomic loci bound by all three features (Fig. 4a). Functional annotation of these sites to genomic compartments revealed a predominant binding of PR-DUB.1 to gene promoters (Fig. 4a), with read densities sharply peaking at TSSs of RefSeq annotated genes (Fig. 4b). Moreover, we found that feature enrichments within ±1kb of TSSs are highly correlated to each other (>0.8), further indicating that ASXL1, FOXK1 and OGT1 are likely subunits of the same protein complex (Fig. 4c). To identify classes of genes bound by PR-DUB.1, we subjected the set of TSSs bound by each complex member to MSigDB pathway enrichment analysis. This analysis identified highly overlapping sets of enriched pathways for each protein (Supplementary Fig. 6b). Notably, PR-DUB.1 targets are predominantly enriched for genes involved in fundamental cellular processes like gene expression, cell cycle, mitosis and protein metabolism (Fig. 4c).

**Figure 4.**
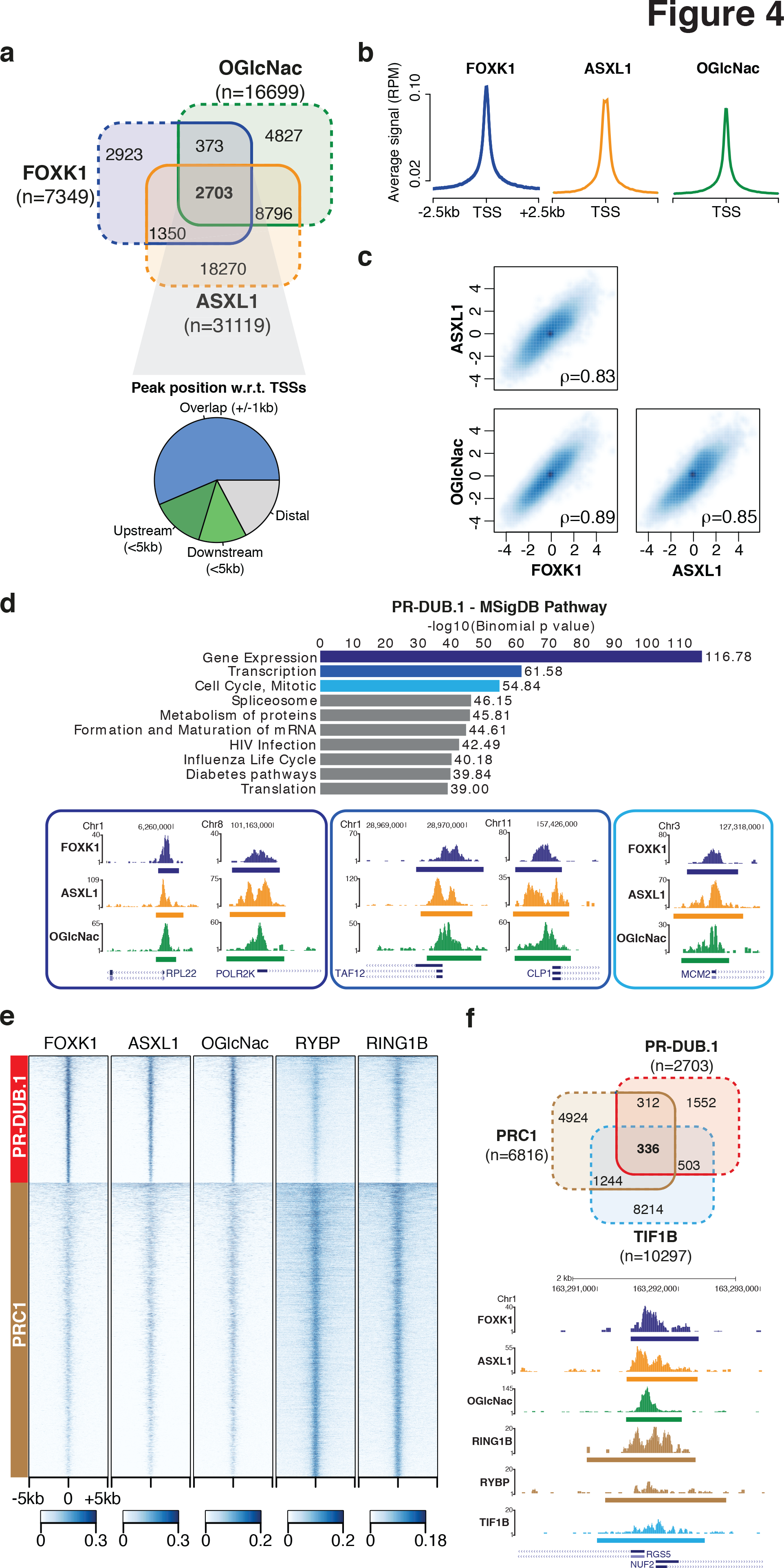
PR-DUB.1 and PRC1 target largely distinct set of genes. (a)Venn diagrams showing the genome-wide colocalization of high-confidence peaks for the PR-DUB.1 components FOXK1 (blue), ASXL1 (orange) and the O-GlcNAc modification (green). Empirical p-values from peak shuffling are indicated, along with the percentage of intersecting peaks for each feature. The pie chart illustrates the distribution of PR-DUB.1 peaks (2703, triple intersection) with respect to TSSs. (b) Average ChIP-seq signal (normalized to total library size) of FOXK1, ASXL1, O-GlcNAc within a 5 kb window centered on RefSeq TSS. (c) Pairwise correlation of PR-DUB.1 feature enrichments at TSSs. Spearman's rank correlation coefficients are indicated. (d) Functional annotation of high-confidence PR-DUB.1 peaks localizing within 5 kb of annotated TSSs. Top 10 significantly enriched MSigDB pathways obtained with GREAT84 are indicated. UCSC tracks of PR-DUB.1 ChIP-seq signals at representative promoters belonging to the top three hits are shown in order of significance (encoded by blue tones). (e) Heatmap of ChIP-seq signals (normalized to total library size) for the indicated features within 10 kb of PR C1 and PR-DUB.1 binding sites. (f) Venn diagrams showing the genome-wide colocalization of high-confidence PR-DUB.1 peaks (red), PRC1 (brown) and TIF1B (light blue). A representative UCSC track of ChIP-Seq signals at TSSs bound by all three features is shown.

### PRC1 complexes and PR-DUB.1 regulate different target genes

Mutations in Drosophila sxc (the gene encoding Ogt), calypso and Asx genes lead to de-repression of HOX genes and previous studies reported a strong colocalization of PR-DUB and O-GlcNAc with major PRC1 bound sites at inactive genes in Drosophila^15, 44^. We sought to investigate this relation in the human genome by comparing our PR-DUB profiles with publicly available ChIP-seq data of RING2 and RYBP^23^, as well as TIF1B^80^.

Our analysis therefore focused on six representatives of the three major modules within our PcG interaction network at the chromatin level: RING2 and RYBP, the central core of the PRC1 module (Fig. 1c); TIF1B, the common component of ZNFs containing CBX1/3/5 complexes (Supplementary Fig. 4a), and PR-DUB.1. Besides the expected high correlation between RING2 and RYBP (p=0.78, Supplementary Fig. 6c), analysis of pairwise correlations of feature enrichments at promoters revealed a clear segregation between PRC1 and TIF1B on the one hand, and PR-DUB.1 on the other hand (Fig. 4e–f and Supplementary Fig. 6c). Similarly, when comparing the genome-wide distribution of PR-DUB.1 (2703 ASXL1+GlcNAc+FOXK1 co-occupied regions) with ?PRC1? (6816 RING2+RYBP peaks) and TIF1B (10297 peaks), we observed only a partial co-localization of these three complexes at target sites, with 24% and 31% of PR-DUB.1 binding sites co-bound by PRC1 and TIF1B, respectively and only 336 regions occupied by all three complexes (Fig. 4f).

In summary, our analysis uncovered the basic topology of the human PR-DUB network at both proteomics and genomics level. Interestingly, and in contrast to Drosophila, the human PR-DUB and PRC1 complexes bind largely distinct sets of target genes, strongly suggesting they are involved in different cellular processes in mammals. In addition, our AP-MS experiments identified the transcription factors FOXK1 and FOXK2 as components of PR-DUB, hence highlighting a potential recruitment mechanism of PR-DUB complexes. We anticipate that future experiments based on our data will shed light on the functionality of PR-DUB complexes in gene regulation and their relation to PRC1 and PRC2.

## Conclusions

Although considerable progress has been made in determining the composition of mammalian PcG protein complexes, recent findings are primarily based on studies of isolated protein components in different cellular contexts with heterogenous biochemical workflows, thus hampering a system-level understanding of gene silencing. In this study, in contrast, we used a systematic proteomic approach to comprehensively map the PcG protein interactome in a single human cell line. Since the abundance of PcG proteins can vary between cell types and surely influences the assembly of alternative protein complexes, we chose HEK293 cells for our study as all PcG proteins are expressed in this cell type. The result is a high-density interaction network, which enabled us to dissect individual PcG complexes with unprecedented detail. By allocating newly identified interaction partners to all PcG complex families and by identifying candidate subunits responsible for complex targeting to chromatin, we obtained new insights into molecular function and recruitment of the PcG silencing system. In addition to the fine mapping of the cardinal PcG complexes PRC1 and PRC2, our data unravel human PR-DUB as multifaceted assembly comprising OGT1 along with several transcription and chromatin binding factors. For the first time, our study testifies the significant diversity that exists among individual PcG complexes in a single cell line. In addition, it provides a solid framework for future systematic experiments aiming at disentangling the biochemistry of PcG protein-mediated gene regulation in mammalian cells.

## Methods

### Expression constructs and generation of stable cell lines

To generate expression vectors for tetracycline-induced expression of N-terminally SH-tagged bait proteins, human ORFs within pDONR223 vectors were picked from a Gateway-compatible human orfeome collection (horfeome v5.1, Open Biosystems) for LR recombination with the customized destination vector pcDNA5/FRT/TO/SH/GW, which was obtained through ligation of the SH-tag coding sequence and the Gateway recombination cassette into the polylinker of pcDNA5/FTR/TO (Invitrogen). Genes not in the human orfeome collection were amplified from human cDNA prepared from HEK293 cells by PCR and cloned into entry vectors by TOPO (pENTR/D-TOPO) reaction. Stable Flp-In HEK293 T-REx cell lines were generated as described in Supplementary Methods.

### Protein purification

Stable Flp-In HEK293 T-REx cell lines were grown in five 14.5 cm Greiner dishes to 80% confluency and bait protein expression induced by the addition of 1*μ*g/ml of tetracycline to the medium 16-24hrs prior to harvest in PBS containing 1 mM EDTA. The suspended cells were pelleted and drained from the supernatant for subsequent shock-freezing in liquid nitrogen and long term storage at −80°C. The frozen cell pellets were resupended in 5ml TNN lysis buffer (100 mM Tris pH 8.0, 5 mM ETDA, 250 mM NaCl, 50 mM NaF, 1% Igepal CA-630 (Nonidet P-40 Substitute), 1.5 mM Na3VO4, 1 mM PMSF, 1mM DTT and 1x Protease Inhibitor mix (Roche)) and rested on ice for 10 min. Insolubilizable material was removed by centrifugation. Cleared lysates were loaded on a pre-equilibrated spin column (Biorad) containing 200 *μ*l Strep-Tactin sepharose (IBA Biotagnology). The sepharose was washed four times with 1 ml TNN lysis buffer (Igepal CA-630 and DTT concentrations adjusted to 0.5% and 0.5mM, respectively). Bound proteins were eluted with 1 ml 2 mM Biotin in TNN lysis buffer (Igepal CA-630 and DTT concentrations adjusted to 0.5% and 0.5mM, respectively), incubated for 2h with 100 yU,l HA-Agarose (Sigma), washed four times with TNN lysis buffer (Igepal CA-630 concentration adjusted to 0.5%, w/o DTT and w/o protease inhibitors) and two additional times in TNN buffer (100 mM Tris pH 8.0, 150 mM NaCl, 50 mM NaF).

The bound proteins were released by acidic elution with 500 *μ*l 0.2 M Glycine pH 2.5 and the eluate was pH neutralized with NH4HCO3. Cysteine bonds were reduced with 5 mM TCEP for 30 min at 37°C and alkylated in 10 mM iodacetamide for 20 min at room temperature in the dark. Samples were digested with 1 pg trypsin (Promega) overnight at 37°C. Bait proteins with low protein yield were processed by single step purification, omitting the HA step. The frozen cell pellets were resuspended in 5ml of TNN lysis buffer containing 10 *μ*g/ml Avidin. The eluates were TCA precipitated to remove biotin and resolubilized in 50 pl 10% ACN, 50 mM NH4HCO3 pH 8.8. After dilution with NH4HCO3 to 5% ACN the samples were reduced, alkylated and digested as in the double step protocol. The digested peptides were puri?ed with C18 microspin columns (The Nest Group Inc.) according to the protocol of the manufacturer, resolved in 0.1% formic acid, 1% acetonitrile for mass spectrometry analysis.

### Mass spectrometry

LC-MS/MS analysis was performed on an LTQ Orbitrap XL mass spectrometer (Thermo Fisher Scientific). Peptide separation was carried out by reverse phase a Proxeon EASY-nLC II liquid chromatography system (Thermo Fisher Scientific). The reverse phase column (75 *μ*m × 10 cm) was packed with Magic C18 AQ (3 *μ*m) resin (WICOM International). A linear gradient from 5% to 35% mobile phase (98% acetonitrile, 0.1% formic acid) was run for 60 min over a stationary phase (0.1% formic acid, 2% acetonitrile) at a ?ow rate of 300 nl/min. Data acquisition was set to obtain one high resolution MS scan in the Orbitrap (60,000 @ 400 m/z) followed by six collision-induced fragmentation (CID) MS/MS fragment ion spectra in the linear trap quadrupole (LTQ). Orbitrap charge state screening was enabled and ions with unassigned or single charge states were rejected. The dynamic exclusion window was set to 15s and limited to 300 entries. The minimal precursor ion count to trigger CID and MS/MS scan was set to 150. The ion accumulation time was set to 500 ms (MS) and 250 ms (MS/MS) using a target setting of 106 (MS) and 104 (MS/MS) ions. After every biological replicate measurement, a peptide reference sample containing 200 fmol of human [Glu1]-Fibrinopeptide B (Sigma-Aldrich) was analyzed to monitor the overall LC-MS/MS systems performance.

### ChIP and preparation of ChIP-seq libraries

Chromatin fixation and immunoprecipitation were performed essentially as described^81^. Cells (3-4×10^8^) were fixed in 200 ml of medium with 1% formaldehyde for 10 min at room temperature. Cross-linked cells were sonicated to produce chromatin fragments of an average size of 150-400 bp. Soluble chromatin was separated from insoluble material by centrifugation. The supernatant containing chromatin of 1-2×10^7^ cells was used for immunoprecipitation. Sequencing libraries were prepared with the NEB Genomic DNA Sample Preparation Kit according to NEB?s instructions. After adapter ligation, library fragments of 250-350 bp were isolated from an agarose gel. The DNA was PCR amplified with Illumina primers with 18 cycles, purified, and loaded on an Illumina flow cell for cluster generation. Libraries were sequenced on the Genome Analyzer IIx (TrueSeq cBot-GA v2 and TruSeq v5 SBS kit) and HiSeq 2000 (HiSeq Flow Cell v3 and TruSeq SBS Kit v3) following the manufacturers protocols. For ChIP-qPCR, nuclei were prepared essentially as described in *Functional Analysis of DNA and Chromatin*^82^. Immunoprecipitations were performed using Anti-GAL4 (sc-510, Santa Cruz Biotechnology), Anti-IgG (10500C, Invitrogen) and Anti-H3K27me3 kindly provided by Thomas Jenuwein. Anti-FOXK1 (ab18196) was purchased from Abcam, Anti-ASXL1 (sc85283) from Santa Cruz Biotechnology and Anti-GlcNAc (HGAC85) from Novus Biologicals. Primer sets used for qPCR are listed in the Supplemental Experimental Procedures.

### Data analysis

Description of data processing and analysis methods are available in the Supplementary Methods.

## Accession Numbers

Mass spectrometry data have been submitted to the PeptideAtlas database http://www.peptideatlas.org/ and assigned the identifier PASS00347. Protein interactions have been submitted to the IMEx (http://www.imexconsortium.org) consortium through IntAct83 and assigned the identifier IM-21659. Sequencing data have been submitted to the NCBI Gene Expression Omnibus (http://www.ncbi.nlm.nih.gov/geo) under accession no. GSE51673.

## Acknowledgements

We thank I. Nissen and M. Kohler for technical support on ChIP-seq. Illumina sequencing was done in the Genomics Facility Basel at D-BSSE, ETH Zurich. Research of SH and MG is supported by the European Union 7th Framework project SYBILLA (Systems Biology of T-cell activation) and the Innovative Medicines Initiative project ULTRA-DD. Research of RA is funded by advanced ERC grant Proteomics v3.0 (233226) and by SystemsX.ch, the Swiss initiative for systems biology. Research of RP is funded by the Swiss National Science Foundation and the ETH Zürich.

## Supplementary Figure Legends

**Supplementary Figure 1.**
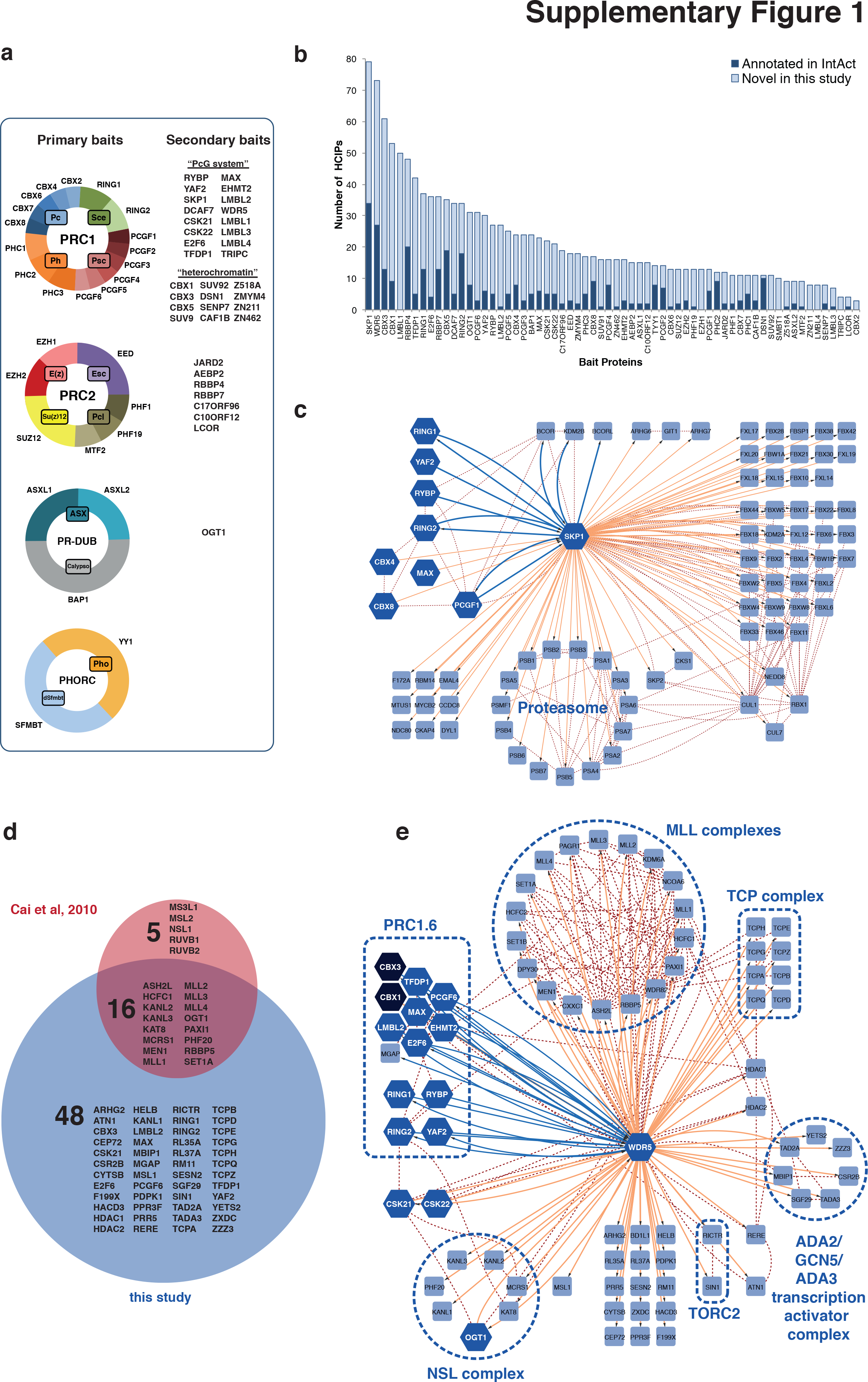
Comparison of AP-MS interaction data with public protein interaction databases and high resolution interaction map of SKP1 and WDR5. (a) Compilation of human bait proteins used for AP-MS and their relation to core subunits of Drosophila PcG complexes. (b) Number of high confidence interacting proteins (HCIPs) for each bait. Dark blue bars represent interactions annotated in the public protein interaction database IntAct, light blue bars are novel interactions found in this study (overall 75%). (c) SKP1 interactome inferred in this study (blue and orange lines) and annotated in public literature databases (red dotted lines). The thick blue lines represent the main interactions to PcG components that cluster together with SKP1 and form PRC1.1. About half of the identified interactions (n = 42) were F-box proteins, including KDM2B – the only F-box protein associated with the PRC1.1 complex. Hexagons indicate bait, squares prey proteins. (d) WDR associated proteins overlapping with^1^. Uniprot protein names and number of proteins are indicated. (e) Interaction map of WDR5 AP-MS results. Blue lines represent interactions detected in this study to PcG associated proteins that cluster with WDR5 and form PRC1.6. Interactions with other HCIPs (orange) together with public literature interactions (dotted red lines) could be assigned to known protein complexes (MLL, TCP, NSL, TORC2 and ADA2/GCN5/ADA3 transcription activator complex). Hexagons indicate bait, squares prey proteins.

**Supplementary Figure 2.**
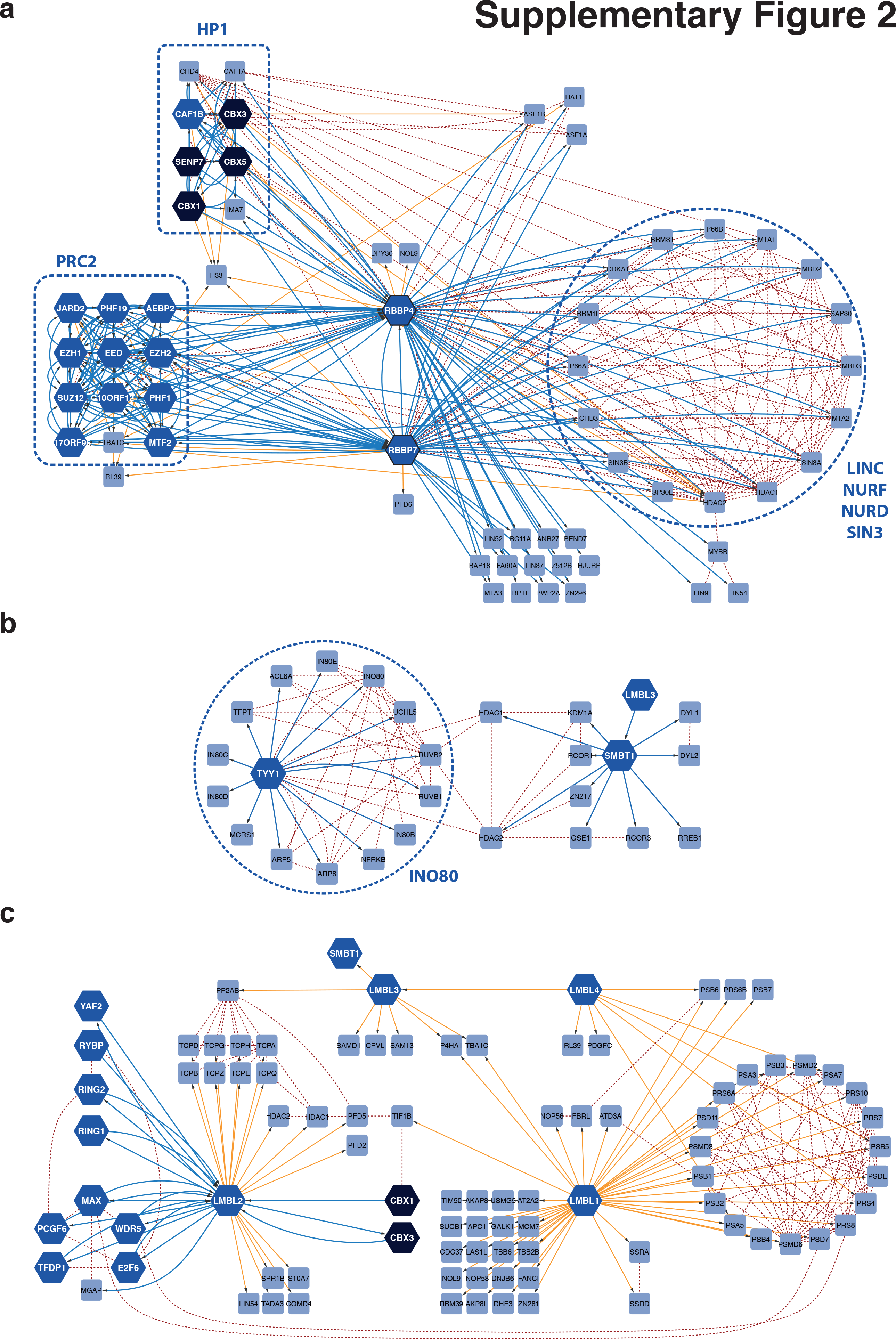
High-resolution topology maps of selected clusters. (a) RBBP4/7 interaction proteome. In addition to interacting with PRC2 and HP1, RBBP4/7 proteins were found to bind members of the LINC, NURF, NURD and SIN3 complexes. (b) No human Pho-RC could be identified. TYY1 and SMBT1, the homologs of the Drosphila Pho-RC proteins Pho and dSFMBT, respectively, did not interact. However, TYY1 co-purified with the INO80 complex, which is indicated by a dashed circle. (c) Interaction proteome of the four human LMBL paralogs. Only LMBL2 exhibited interactions with a PcG protein assembly and is part of PRC1.6. In all panels blue and orange lines represent interactions found in this study; red dashed lines are annotated public literature interactions. Hexagons indicate bait, squares prey proteins.

**Supplementary Figure 3.**
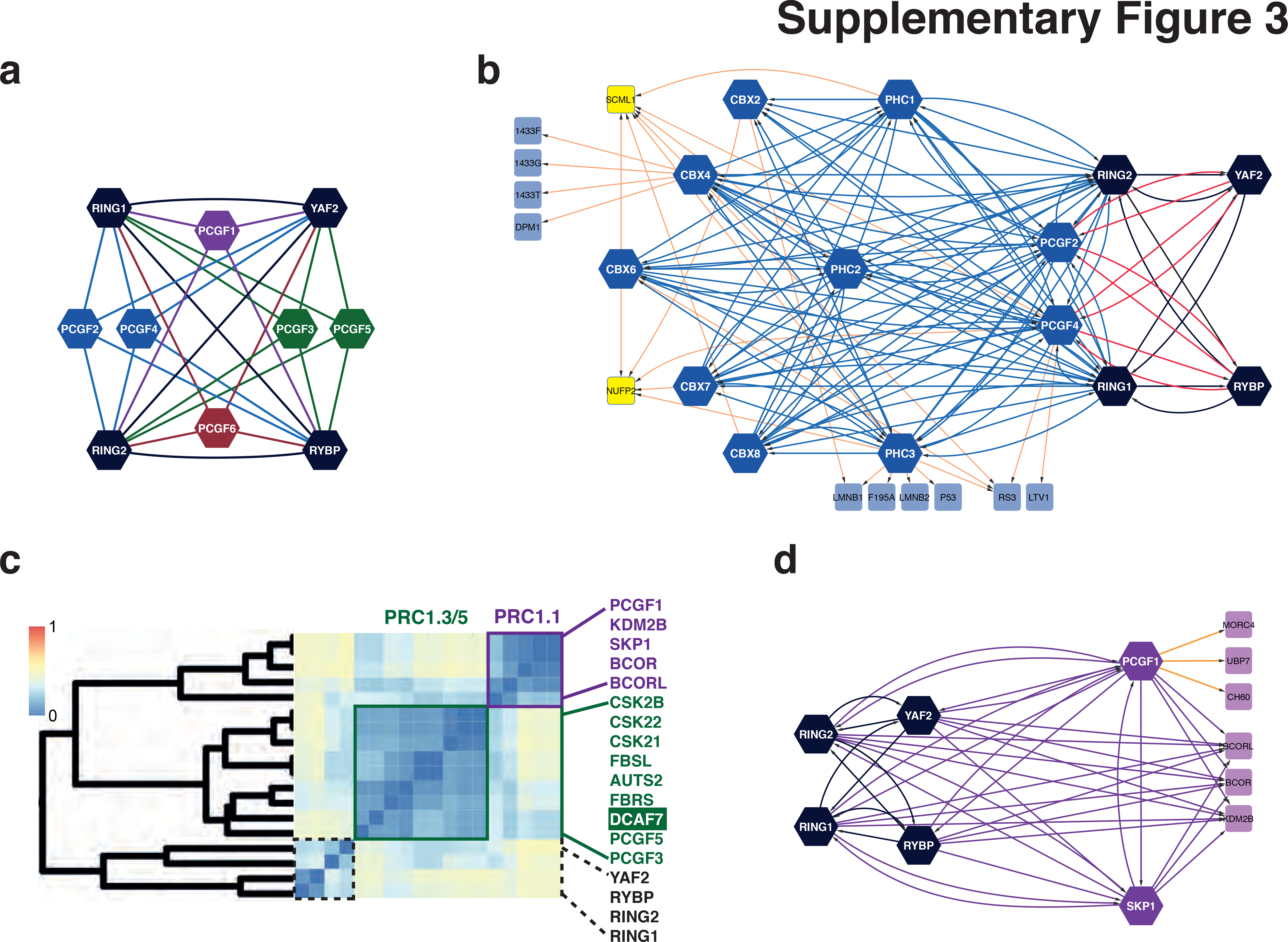
Network topology of PRC1 complexes. (a) Protein-protein interactions among the central core proteins of the PRC1 complexes. Note that we detected no interactions between protein paralogs. Edges represent detected reciprocal interactions. (b) Network topology of PRC1.2 and PRC1.4 complexes. Note that PCGF2/4 can assemble to PRC1.2/PRC1.4 or can form trimeric complexes (edges in red) with RING1/2 and RYBP/YAF2 (we detect no interactions between RYBP/YAF2 with CBX and PHC proteins). Edges connecting PRC1.2/PRC1.4 core components in blue. (c) Excerpt of Figure 2A indicating the allocation of the WD40 protein DCAF7 to PRC1.3/PRC1.5. High-density interaction maps of PRC1.3/PRC1.5 (C) andPRC1.6 (D). New subunits are highlighted by dashed boxes. Hexagon shaped nodes represent baits; squares: identified HCIPs not used as baits in this study. Black nodes: common core subunits; yellow nodes: DNA binding proteins. (d) PRC1.1 complex contains PCGF1 and SKP1 and links the co-repressors BCOR and BCORL as well as the demethylase KDM2B to the PRC1.1 core. Edges connecting core components are in purple.

**Supplementary Figure 4.**
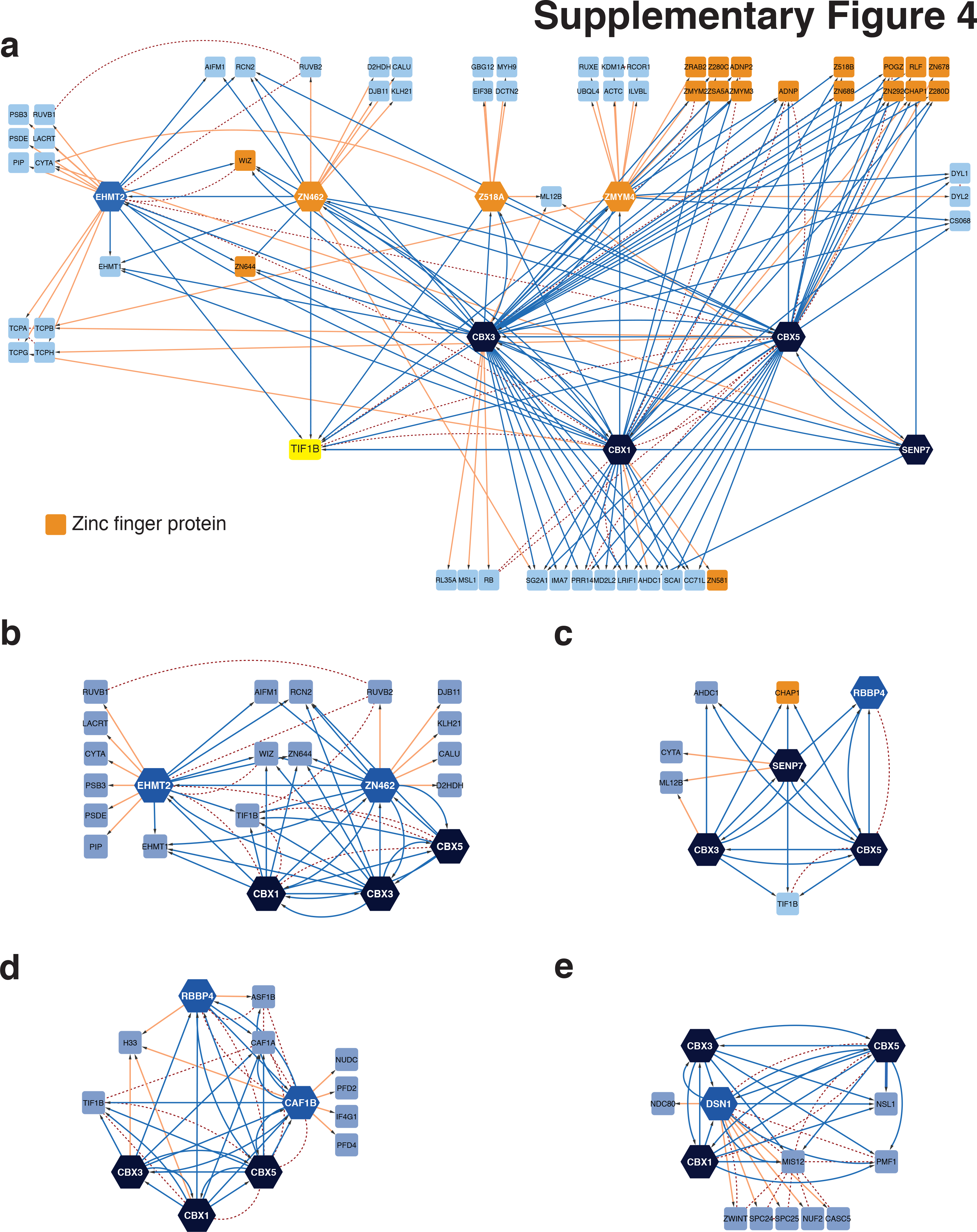
Interaction proteomes of PRC1 and HP1 complexes. (a) CBX1/3/5 interactions with zinc finger (ZNF; orange nodes) and neighboring proteins. Note that all bait proteins of this subnetwork interacted with the nuclear co-repressor TIF1B (yellow), which underscores the reported function of TIF1B in the recruitment of HP1 proteins to specific DNA sequences through interaction with zinc finger transcription factors. Hexagons indicate bait, squares prey proteins. Blue edges, interactions of potential core subunits of the corresponding network, interactions in public databases as dashed lines, others in orange. (b) CBX1 and CBX3 but not CBX5 interact with the histone methyltransferases EHMT1 and EHMT2. The CBX1/3-EHMT1/2 complex also interacts with zinc finger transcription factors WIZ, ZN644, ZN462 and the co-repressor protein TIF1B. In contrast to a previous report we did not detect any potential interactions of EHMT1/2 with PRC1.6^2^. (c) Interaction of CBX3 and CBX5 with SENP7. This complex may also include TIF1B, the zinc finger transcription factor AHDC1 and the histone chaperone CHAP1. (d) CBX1/3/5 complex with histone chaperone CAF1 and RBBP4, which is potentially involved in DNA replication. (e) Centromeric DSN1/MIS12 complex with HP1 proteins, involved in mitosis.

**Supplementary Figure 5.**
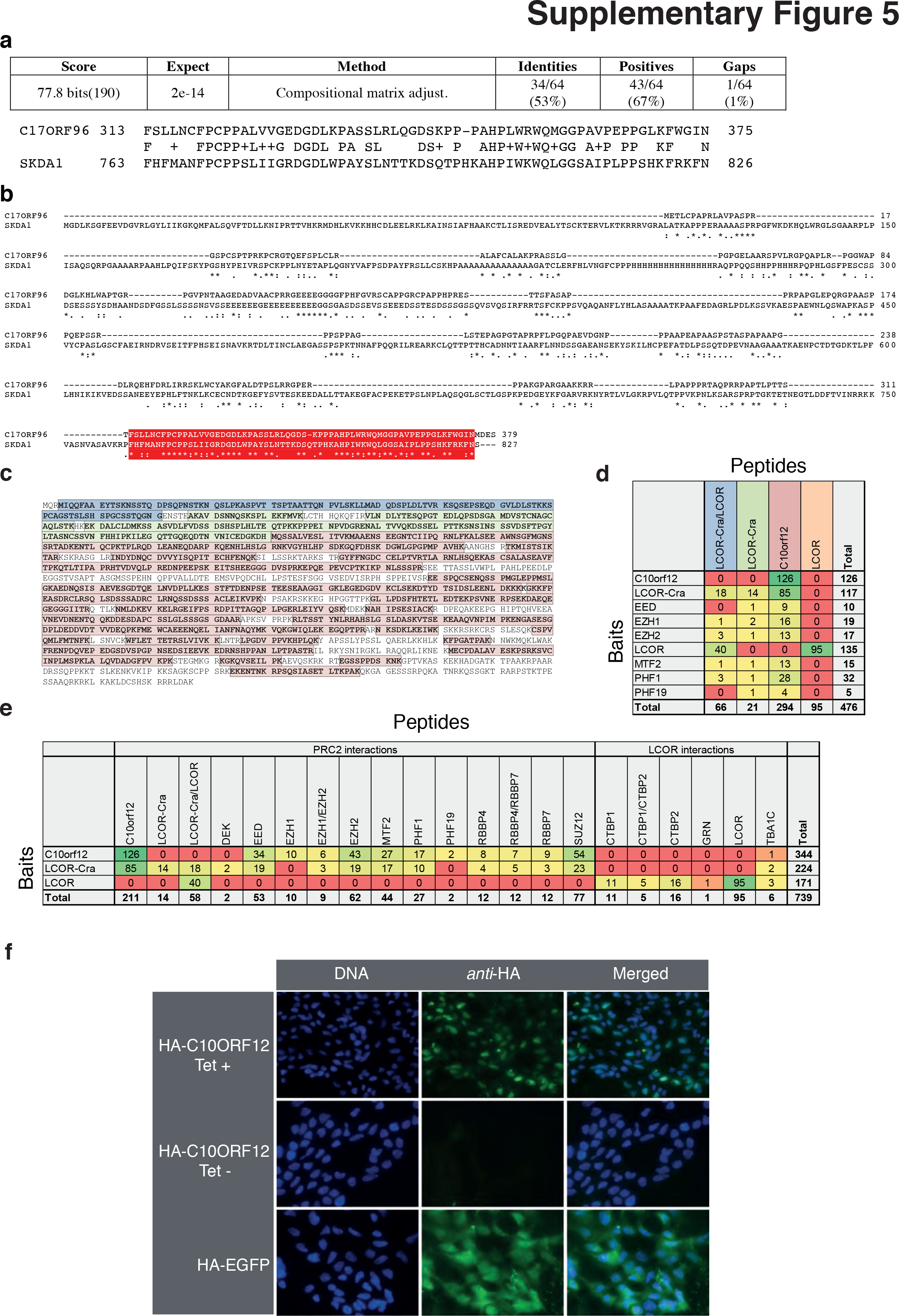
Characterization of PRC2 core interacting proteins C17ORF96 and C10ORF12. (a) C17ORF96 and SKDA1 are related proteins, which share sequence similarity at their C-terminus. BLAST search result with C17ORF96 (Uniprot A6NHQ4) as query sequence. (b) CLUSTAL 2.1 alignment of C17ORF96 (Uniprot A6NHQ4) and SKDA1 (Uniprot Q1XH10) amino acid sequences. Homologous C-termini, identified by BLAST search, indicated in red. (c) All AP-MS identified peptides that match to LCOR-CRA_b. Colors indicate specific protein regions as in Fig. 2c. (d) Number of identified LCOR and C10ORF12 isoform peptides in PRC2 protein purification experiments. (e) Number of PRC2 prey peptides in LCOR and C10ORF12 isoform AP-MS experiments. (f) C10ORF12 localize in the nucleus. Images show anti-HA in situ stainings of stable Flp-In HEK293 T-REx cell lines before and after tetracycline induction. HA-EGFP shows a dispersed, cytoplasmic signal whereas the C10ORF12 HA-epitope fusion protein shows a nuclear signal. In situ stainings were performed as described in Glatter et al., 2009^3^.

**Supplementary Figure 6.**
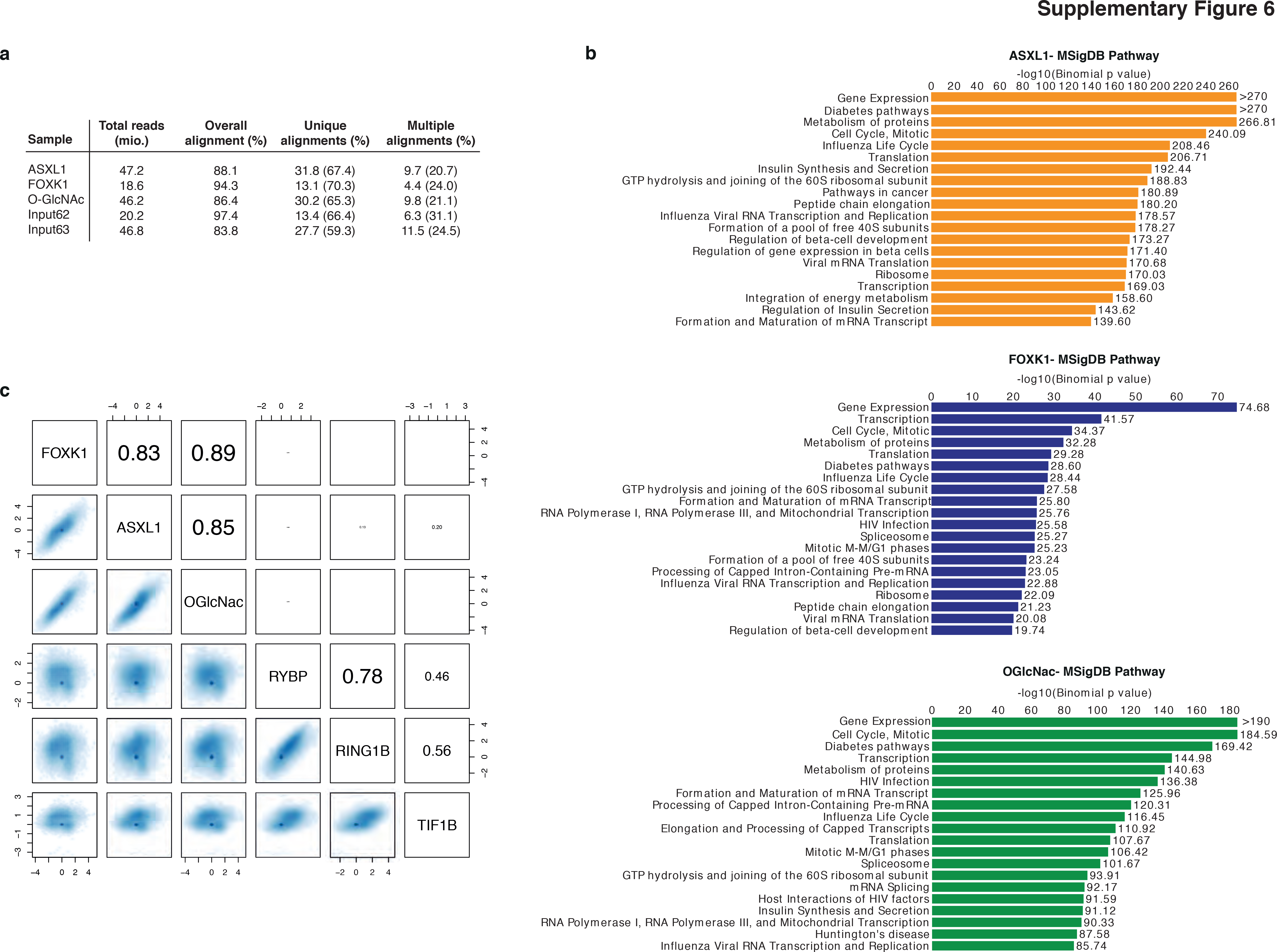
Functional annotation of PR-DUB.1 components binding sites and genome-wide analysis of PR-DUB.1, PRC1 and TIF1B enrichments at TSSs. (a) Sequence read statistics of ChIP-seq experiments. (b) Functional annotation of high-confidence MACS peaks localizing within 5kb of annotated TSSs. Enriched MSigDB pathways were computed with GREAT^4^. (c) Pairwise scatter plots of PR-DUB.1, PRC1 and TIF1B enrichments at TSSs. Spearman’s rank correlation coefficients are indicated.

## Supplementary Methods

### Cell Line Generation

Flp-In HEK293 T-REx cells (Invitrogen) containing a single genomic FRT site and stably expressing the tet repressor were cultured in DMEM (4.5 g/l glucose, 10% FCS, 2 mM L-glutamine) containing 100 *μ*g/ml zeocin and 15 *μ*g/ml blasticidin. The medium was exchanged with DMEM medium containing 15 *μ*g/ml blasticidin before transfection. For cell line generation, Flp-In HEK293 T-REx cells were co-transfected with the corresponding expression plasmids and the pOG44 vector (Invitrogen) for co-expression of the Flp-recombinase using the Lipofectamine 2000 transfection reagent (Invitrogen). Two days after transfection, cells were selected in hygromycin-containing medium (100 *μ*g/ml) for 2-3 weeks.

### Protein Identification

Mass spectrometry raw data were searched with X!Tandem^5^ against a human protein sequence database (Swiss-Prot canonical reviewed human proteome reference data set; http://www.uniprot.org/), including reverse decoy sequences for all entries. The search parameters were set to include only fully tryptic peptides (KR/P) containing up to two missed cleavages. Peptide modifications consisted of Carbamidomethyl (+57.021465 amu) on Cys (static) and oxidation (+15.99492 amu) on Met (dynamic) and phosphorylation (+79.966331 amu) on Ser, Thr, Tyr (dynamic) were set as dynamic peptide modifications. Precursor mass error tolerance was set to 25 ppm, the fragment mass error tolerance to 0.5 Da. Obtained peptide spectrum matches were statistically evaluated using PeptideProphet and protein inference by ProteinProphet, both part of the Trans Proteomic Pipeline^6^. A minimum protein probability of 0.9 was set to match a false discovery rate (FDR) of <1%. The resulting pep.xml and prot.xml files were used as input for the spectral counting software tool Abacus7 to calculate spectral counts and normalized spectral abundance factor (NSAF) values^8, 9^.

### Evaluation of high confidence interacting proteins (HCIP)

Adjusted NSAF values of identified co-purified proteins were compared to a control data set of 62 StrepHA-GFP and 12 StrepHA-RFP-NLS purification experiments. The protein abundance in the control data set was estimated by averaging the 10 highest NSAF values per protein among all 74 measurements. Protein abundance enrichment of >10 fold compared to the control data set was used as an initial step for filtering protein interaction raw data. Adjusted NSAF values were also used to calculate WDN-scores of all the interaction candidates^10^. A simulated data matrix was used to calculate the WD-score threshold below which 98% of the simulated data falls. From this high confidence interaction data set (control ratio > 10; WDN-score > 1) a distance matrix was calculated with the Multiple Experiment Viewer^11^ (http://www.tm4.org/mev/) using an uncentered Pearson distance metric and mapped on the unfiltered raw interactions. To relax filtering stringency in close proximity in the network, sub-threshold interactions (control ratio and WDN-score) were rescued if the distance was greater than zero (n = 314 protein interactions). The resulting filtered data set contained the high confidence interacting proteins (HCIPs) and corresponding protein-protein interactions. For comparison to literature data, all human protein interactions were extracted from the public database IntAct^12^.

### Clustering analysis

All data analyses were performed using R^13^ (http://www.R-project.org). Agglom-erative hierarchical clustering of HCIP was performed using adjusted NSAF values. Different correlation-based dissimilarity measures were considered in combination with commonly adopted intergroup dissimilarity measures (single, average and complete linkage functions). For each pair of measures, clustering performances were evaluated using the cophenetic correlation coefficient, which measures the ability of a dendrogram to represent the input data structure^14^. As a result of this procedure, hierarchical clustering was performed by adopting a Spearman’s rank correlation coefficient based dissimilarity along with average linkage. Therefore, the dissimilarity between prey i and j was computed as *d_ij_* = (1 − *r*(*x_i_*, *X_j_*))/2, where *r* is the SCC.

### Network Visualization

Protein Interaction data were visualized with Cytoscape 2.8.3^15^. Known bait interactions were obtained from the protein interaction network analysis platform PINA v2 (December 2012)^16^ using bait protein identifiers as starting nodes.

### ChIP-Seq data analysis

ChIP-Seq profiles of RING1B, RYBP, TIF1B (GEO accession number GSM855007, GSM855008 and GSE27929, respectively) and corresponding input data sets were downloaded in sra format and converted to fastq using the NCBI Short Read Archive Toolkit. Short reads were aligned to the human genome (hg19 assembly) using Bowtie 2.0.0^17^ allowing for 1 mismatch in a 30nt seed, reporting best out of at most 100 alignments. Overall alignment rates ranged between 83 and 94% for in-house generated data sets and between 75 and 98% for the others. Alignments were converted from SAM format to BAM using SAMtools 0.1.18^18^. Peak calling was performed using MACS 1.4.0^19^ with default parameters. Peaks were then filtered according to p-values (p < 10^−10^). If replicates were available, only peak intersections were considered further and denoted as high-confidence peaks. All subsequent analyses were performed using R/Bioconductor^20^. Coverage tracks at single base pair resolution were generated with wavClusteR^21^. Overlapping peaks were determined using GenomicRanges using a minimum overlap of 1bp. RefSeq transcript annotations were fetched from UCSC using GenomicFeatures. Unique TSSs were defined as TSSs having no other annotated TSS within their 1kb flanking region, irrespective of the strand. A total of 21612 unique TSSs was considered further. Metaprofiles of ChIP-Seq signals at TSSs (± 2.5kb) were computed using non-overlapping windows of width 50 nt. ChIP-Seq signal heatmaps were computed with Genomation^22^. Feature enrichments at unique promoters were computed as described in^23^. MSigDB pathway analysis was performed with GREAT 2.0.2^4^ by associating genomic regions to single nearest annotated genes within 5kb.

